# Facial shape and allometry quantitative trait locus intervals in the Diversity Outbred mouse are enriched for known skeletal and facial development genes

**DOI:** 10.1101/787291

**Authors:** David C. Katz, J. David Aponte, Wei Liu, Rebecca M. Green, Jessica M. Mayeux, K. Michael Pollard, Daniel Pomp, Steven C. Munger, Steven A. Murray, Charles C. Roseman, Christopher J. Percival, James Cheverud, Ralph S. Marcucio, Benedikt Hallgrímsson

**Affiliations:** Department of Cell Biology & Anatomy, Alberta Children’s Hospital Research Institute and McCaig Bone and Joint Institute, Cumming School of Medicine, University of Calgary, AB, Canada; Department of Molecular Medicine, The Scripps Research Institute, La Jolla, CA, USA; Department of Genetics, University of North Carolina Medical School, Chapel Hill, NC, USA; The Jackson Laboratory, Bar Harbor, ME, USA; Department of Animal Biology, University of Illinois Urbana Champaign, Urbana, IL, USA; Department of Anthropology, Stony Brook University, Stony Brook, NY, USA; Department of Biology, Loyola University Chicago, Chicago, IL, USA; Department of Orthopaedic Surgery, School of Medicine, University of California San Francisco, San Francisco, CA, USA

## Abstract

The biology of how faces are built and come to differ from one another is complex. Discovering the genes that contribute to differences in facial morphology is one key to untangling this complexity, with important implications for medicine and evolutionary biology. This study maps quantitative trait loci (QTL) for skeletal facial shape using Diversity Outbred (DO) mice. The DO is a randomly outcrossed population with high heterozygosity that captures the allelic diversity of eight inbred mouse lines from three subspecies. The study uses a sample of 1147 DO animals (the largest sample yet employed for a shape QTL study in mouse), each characterized by 22 three-dimensional landmarks, 56,885 autosomal and X-chromosome markers, and sex and age classifiers. We identified 37 facial shape QTL across 20 shape principal components (PCs) using a mixed effects regression that accounts for kinship among observations. The QTL include some previously identified intervals as well as new regions that expand the list of potential targets for future experimental study. Three QTL characterized shape associations with size (allometry). Median support interval size was 3.5 Mb. Narrowing additional analysis to QTL for the five largest magnitude shape PCs, we found significant overrepresentation of genes with known roles in growth, skeletal development, and sensory organ development. For most intervals, one or more of these genes lies within 0.25 Mb of the QTL’s peak. QTL effect sizes were small, with none explaining more than 0.5% of facial shape variation. Thus, our results are consistent with a model of facial diversity that is influenced by key genes in skeletal and facial development and, simultaneously, is highly polygenic.

**Author Summary:** The mammalian face is a complex structure serving many functions. We studied the genetic basis for facial skeletal diversity in a large sample of mice from an experimental population designed for the study of complex traits. We quantified the contribution of genetic variation to variation in three-dimensional facial shape across more than 55,000 genetic markers spread throughout the mouse genome. We found 37 genetic regions which are very likely to contribute to differences in facial shape. We then conducted a more detailed analysis of the genetic regions associated with the most variable aspects of facial shape. For these regions, a disproportionately large number of genes are known to be important to growth and to skeletal and facial development. The magnitude of these genetic contributions to differences in facial shape are consistently small. Our results therefore support the notion that facial skeletal diversity is influenced by many genes of small effect, but that some of these small effects may be related to genes that are fundamental to skeletal and facial development.

## Introduction

The facial skeleton is a complex morphology that serves a diverse set of critical roles. It houses and protects the forebrain and most sensory organs and provides essential anatomy for oral communication, feeding, and facial expression. Structured differences (covariance) in facial shape and form depend upon variation in the biological processes that coordinate facial development [1–7]. Understanding how these processes act and combine to build a facial skeleton requires interrogation at multiple levels of the genotype-phenotype (GP) map [8, 9].

Among the most fundamental of these inquiries is the search for genetic variants that contribute to facial variation. These genetic variants are the basic building blocks of diversity and, within species, circumscribe the potential for short-term evolutionary change [10, 11]. Over the past decade, dozens of studies in humans, mouse models, and other species have dissected GP associations for the craniofacial complex using moderate-to-high density genetic maps (tens of thousands to millions of markers). The focus is typically on additive effects [12], though some recent work considers interactions among loci [4, 13, 14].

In humans, genome-wide association studies (GWAS) targeting variation in soft tissue facial form have been completed for European [15–20], African [21], South American [22], and Asian [23] cohorts. Collectively, the studies report around 50 loci for facial morphology [24]. While there is limited overlap in the candidate gene sets among studies [25], and though many of the individual genes have no previously known role in facial development [3], the genes that do replicate across GWAS (*PAX 1, PAX3, HOXD* cluster genes, *DCHS2, SOX9*) are known to be important to facial and/or skeletal morphogenesis.

Compared to human studies, mapping the genetics of facial variation in mice offers several benefits, including greater control over genetic and environmental variation and the ability to record observations, directly or *in silico*, on the skeleton itself [14, 26–31]. In recent years, the focus of mouse skull QTL analyses has expanded to include tests of enrichment for disease phenotypes and roles in skull development, as well as the search for and validation of candidate genes [32–34]. Dozens of QTL have been identified. As with human facial shape GWAS candidates, this is encouraging but also a small beginning—on the order of the total number of loci identified for human height just a decade ago [35]. Quantitative genetics analyses [32, 36, 37] and expression and epigenomic atlases of facial development both strongly point to the potential for hundreds or thousands of genetic loci with as-yet-undetected QTL variance contributions [38–40]. Increased sample sizes, greater mapping resolution, and the development of new experimental lines all have the potential to expand the facial shape QTL library and increase the precision of variant localization [41–43].

The present study maps mouse facial shape QTL in what is, to date, the largest sample yet used for this purpose (*n* = 1147). We use Diversity Outbred mice (DO; Jackson Laboratory, Bar Harbor, ME), a random outcross model derived from the Collaborative Cross (CC) founding lines [43–46]. The DO is specifically designed for genetic mapping of complex traits. Each animal’s genome is a unique mosaic of the genetic diversity present in the CC founders—more than 45 million segregating SNPs [47]. Random outcrossing (of non-sibs) over many DO generations maintains this diversity and increases recombination frequency, which is critical for reducing confidence intervals of localized QTL [41]. Compared to classic backcross and intercross designs, the combinatorial possibilities for allelic variation within and among DO animals are vast. The genetic diversity in a DO sample may thus mimic diversity in the natural populations that animal models are meant to illuminate.

We quantify facial shape with 22 three-dimensional landmarks **(Fig 1)** and estimate its association with founder haplotype variation at 56,885 genetic markers. The dependent observations in the QTL scans are scores on shape PCs. For each PC explaining more than 1% of shape variation, we identify statistically significant marker effects using a mixed effects regression model that simultaneously accounts for kinship among animals [48, 49]. We then undertake a more detailed analysis of QTL intervals for shape PCs 1-5. We estimate founder coefficients, test whether the overall gene set is enriched for biological processes relevant to facial development and perform association mapping to highlight a list of potential causal genes. In order to visualize and quantify the magnitude of marker effects on multivariate facial shape (as opposed to univariate PC scores), for each QTL peak, we fit an additional mixed model that accommodates high-dimensional dependent observations [50].

**Fig 1.**
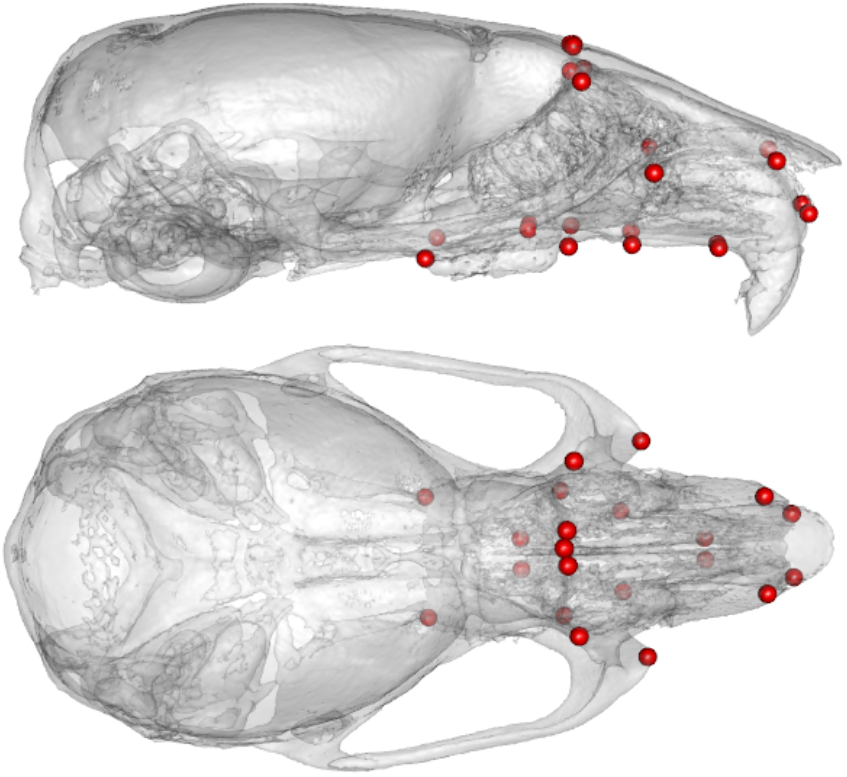
Landmark map.

## Results and Discussion

### QTL and Shape Variable Heritability

We used a mixed effects regression model to quantify GP associations between 56,885 genetic markers and each of the 20 shape PCs explaining >1% of the symmetric component of facial shape variation **(S1 Fig)**. Evidence for a QTL was evaluated by comparing logs odds (LOD) scores (likelihood ratios comparing the magnitude of marker model residuals to the residuals of a null model that excludes the marker effect) [51] at each marker to a distribution of maximum LOD scores obtained from quantifying marker effects over 1000 random permutations of GP assignments.

While comparisons to prior studies are complicated by design features unique to each, the design employed here appears to have conferred benefits in terms of QTL identification and localization, the primary objectives of QTL analysis [41]. From the 20 genome scans, 37 QTL surpassed our threshold for statistical significance (95% quantile of the maximum LOD score distribution). Median support interval width was 3.5 Mb (mean 7.1 Mb) using a 1.5 LOD-drop threshold to define support interval boundaries. Comparable prior studies identified 17-30 QTL for the skull, with a lowest median QTL interval of 7.8 Mb (using a less-demanding 1.0 LOD-drop threshold) [32, 33].

**Fig 2** plots the support intervals for these QTL (**see also S1 Table**). The QTL span 15 of 19 autosomes as well as the X-chromosome. On chromosomes 7, 14, 15, and 17, QTL from multiple PCs have overlapping associations with the same genomic region.

**Fig 2.**
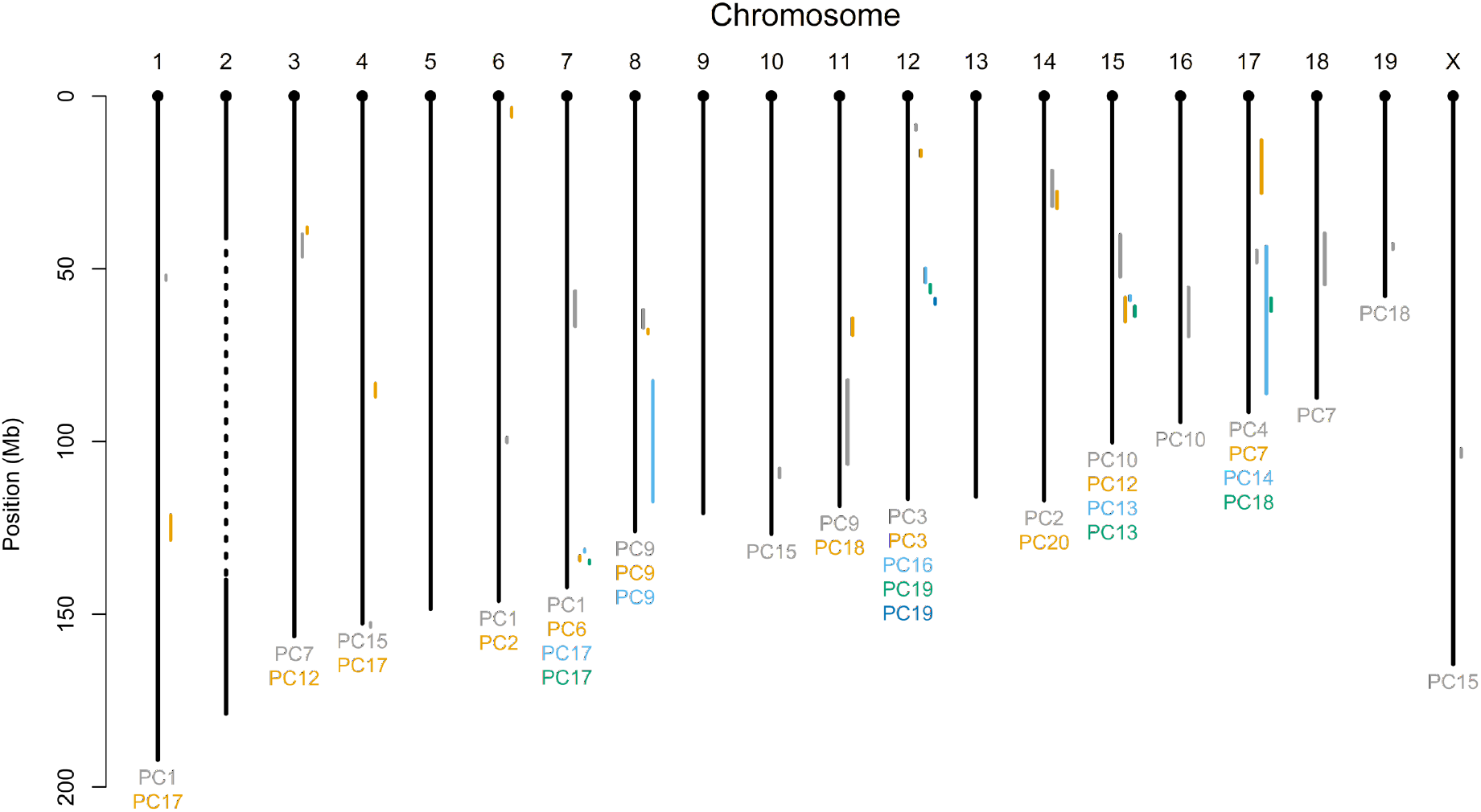
Mouse genome physical map with QTL support intervals. Intervals on the same chromosome are color coded to match the color of the PC for which the QTL was identified (listed below chromosome). The dashed line in the middle of chromosome 2 represents a region for which genetic effects were not evaluated due to the potential effects of transmission distortion on QTL identification (see Materials & Methods for further explanation).

For each PC, the random effect heritability estimate quantifies the proportion of phenotypic variance attributable to overall genetic similarity at the founder level [49]. We excluded from the genetic similarity (relatedness) calculation all markers on the same chromosome as the QTL marker of interest. This leave-one-chromosome-out (LOCO) strategy was chosen in order to avoid the loss of power that can arise when relatedness is computed from the full complement of genotyped markers [52–54]. The LOCO approach produces a unique heritability estimate for each chromosome on each PC. **Table 1** reports the mean heritability for each PC, as well as the variance in these heritabilities (averaged over chromosomes). For most PCs, heritability was moderately high (0.42-0.57), with a few low heritability outliers. These values are very similar to those Pallares and colleagues estimated for skull shape PCs in an outbred mouse sample [32], and lie mostly in the middle of the heritability range for human facial shape (∼30-70%) [36, 55].

**Table 1.** Heritabilities by PC.

|  | <u>Mean $h^2$</u> | <u>Variance</u> |
| --- | --- | --- |
| PC1 | 0.5407 | 0.00012 |
| PC2 | 0.4389 | 0.00013 |
| PC3 | 0.5418 | 0.00022 |
| PC4 | 0.3532 | 0.00022 |
| PC5 | 0.4627 | 0.00012 |
| PC6 | 0.5705 | 0.00015 |
| PC7 | 0.4785 | 0.00014 |
| PC8 | 0.4604 | 0.00015 |
| PC9 | 0.5411 | 0.00032 |
| PC10 | 0.5420 | 0.00017 |
| PC11 | 0.2439 | 0.00009 |
| PC12 | 0.4226 | 0.00014 |
| PC13 | 0.5309 | 0.00013 |
| PC14 | 0.4364 | 0.00020 |
| PC15 | 0.4849 | 0.00005 |
| PC16 | 0.4456 | 0.00010 |
| PC17 | 0.4658 | 0.00020 |
| PC18 | 0.4726 | 0.00019 |
| PC19 | 0.4533 | 0.00006 |
| PC20 | 0.5040 | 0.00022 |

Mean and variance of heritability for PC 1-20 shape traits. The full set of heritability estimates is provided in **S2 Table**.

Based on signal-to-noise evaluation of shape PC eigenvalue decay [56], we limited further analysis to the first five PCs (effectively four PCs, as no marker exceeded the permutation significance threshold for the PC 5 shape axis). Each plot in **Fig 3** depicts the LOD score profile (lower panel) and CC founder haplotype coefficients (upper panel) for a chromosome harboring one or more QTL for a PC 1-4 shape variable. Shaded rectangular regions delimit the 1.5-LOD drop support interval boundaries for the QTL. Founder coefficients were estimated using best linear unbiased predictors [57, 58]. This helps to highlight QTL regions by imposing a degree of shrinkage on the coefficients chromosome-wide [48].

**Fig 3.**
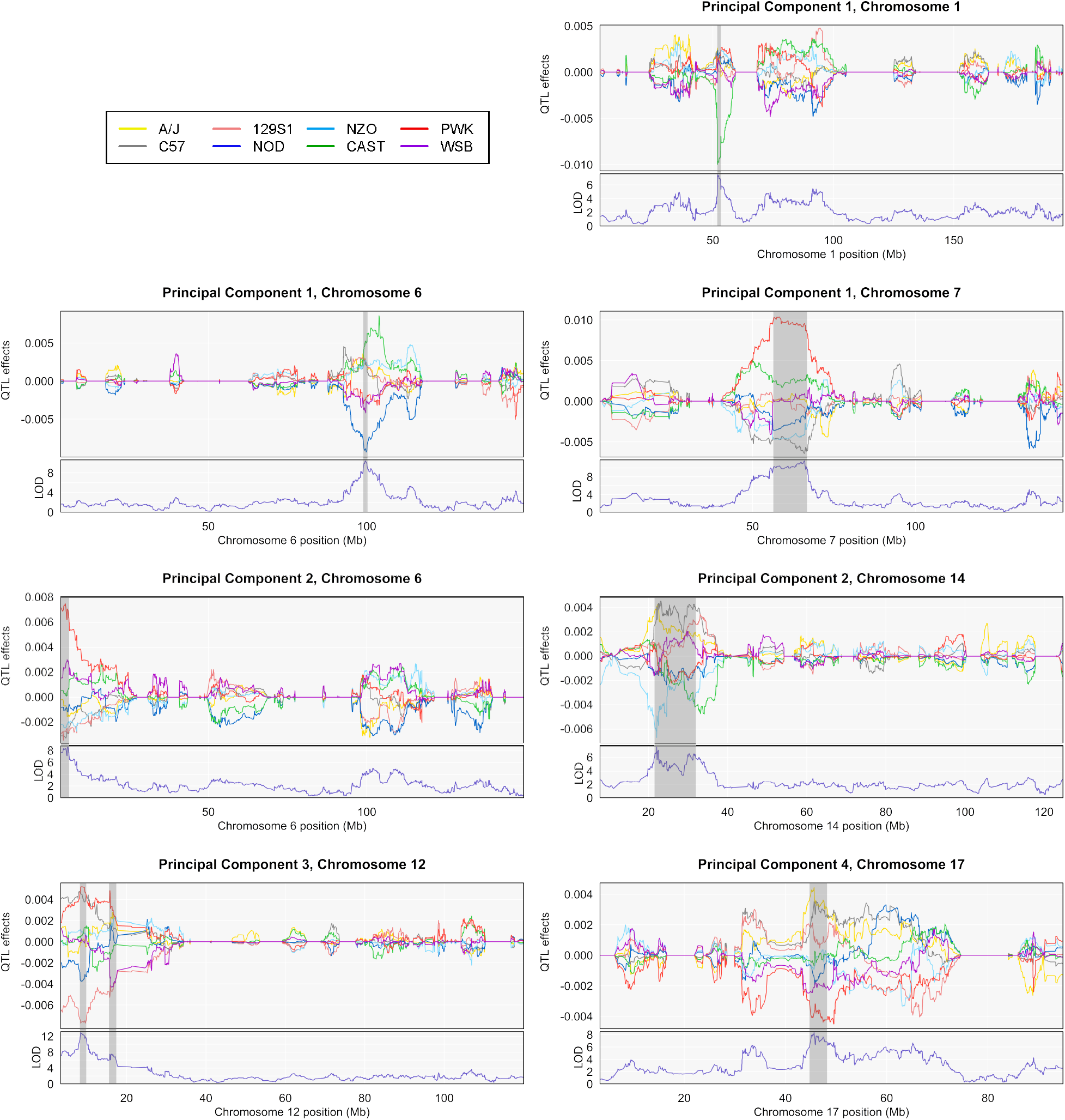
BLUP effects and LOD panels for all QTL, PCs 1-4. QTL effect colors correspond to CC founders.

We also mapped GP associations for vectors capturing covariation between shape and two measures of size: centroid size (CS) and body mass. Shape change associated CS and body mass variation were closely correlated with the primary axis of shape variation in our sample (*r*_PC1-CS_: 0.985, *r*_PC1-Mass_: 0.875, *r*_CS-Mass_: 0.927). These correlations carry forward to the QTL profiles, as well. We identify two QTL each for CS and body mass allometry, one of which, on chromosome 6, is common to the two allometry vectors. Together, the three QTL—on chromosomes 1, 6, and 7—completely overlap with the PC 1 shape QTL (**Fig 4**).

**Fig 4.**
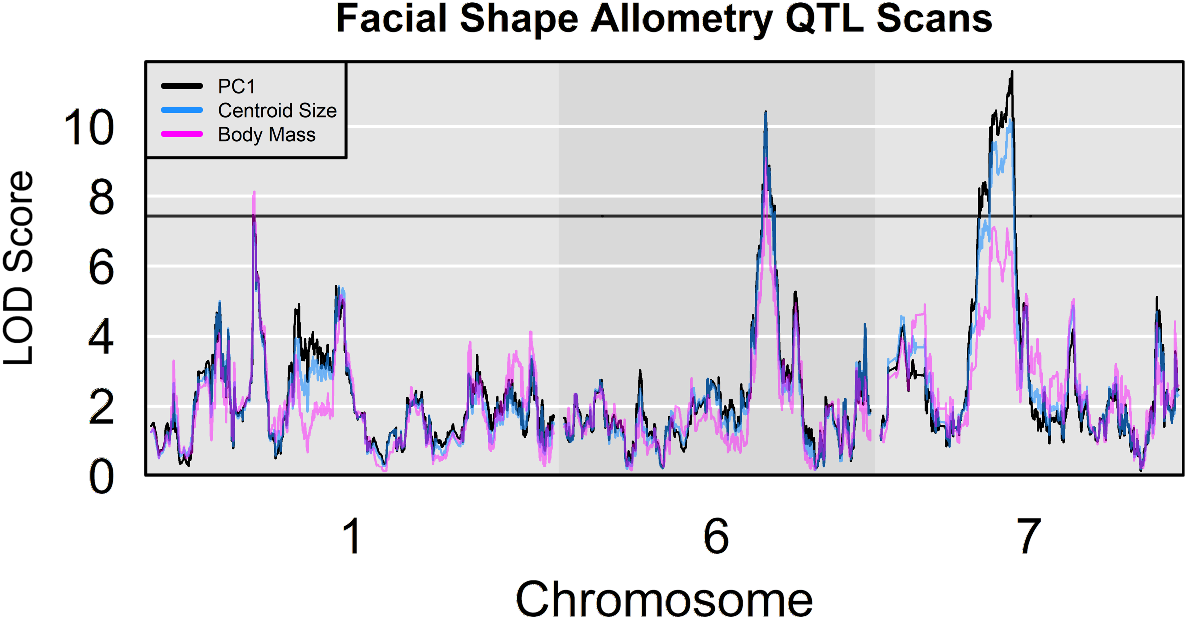
LOD plots for chromosomes containing allometry (CS and body mass) and PC 1 QTL. The permutation significance thresholds for all three phenotypic variables are extremely similar. Only one is plotted.

The allometry results suggest a developmental model in which the determinants of different size measures have partially correlated effects on cranial shape [59]. The near coincidence of shape PC 1 with our explicitly allometric variables has been observed in many geometric and classical morphometric investigations, and there is a long history of treating PC 1 as an allometry variable in morphometric analysis [59–64]. With this in mind, though primarily for economy, we regard PC 1 QTL as allometry QTL in the remainder of this manuscript.

The mouse subspecies that comprise the CC are estimated to have diverged several hundred thousand years ago [65]. In many DO studies, subspecific divergence in one or more wild-derived CC lines figures prominently in the founder genetic effects contrasts that underlie QTL (e.g., [66–69]). Here, the same pattern is evident for most of the PC 1-4 QTL. In six of the eight QTL (or seven of nine, if the two peaks of the PC 2, chromosome 14 QTL are considered distinct), the coefficient for a non-*domesticus* wild-derived strain (CAST or PWK) sits at one extreme of the founder coefficient distribution. Only at the upstream LOD peak of the PC 2, chromosome 14 QTL is the founder coefficient contrast clearly between *M.m. domesticus* inbred lines. The prevalence of QTL substantially driven by wild-derived alleles highlights the value of subspecific genetic diversity in the DO population.

The broad region of high LOD scores for PC 1 on chromosome 7 merits individual attention. This QTL support interval essentially delimits the mouse homolog boundaries of the human Prader-Willi/Angelman syndromes domain [70]. The founder coefficients point to a strong subspecies signal, with PWK (*M.m. musculus*) highly differentiated from wild-derived WSB and the inbred *M.m. domesticus* lines, and CAST (*M.m. castaneus*) intermediate. Though the QTL interval contains fewer than 40 genes, nine are known to be imprinted [70–74]. Several more have mixed evidence for imprinting [73], and Prader-Willi and Angelman are parent-of-origin dependent neurological disorders [75]. In addition, more than a dozen genes in the interval are expressed in the developing face [40, 76–81]. Our mapping strategy is not specifically designed to quantify imprinted gene QTL [82–84]. However, closer dissection of this interval’s contribution to facial shape variation, accounting for allelic parent of origin effects, may be warranted in the future. The characteristic facial phenotype for Prader-Willi includes a relatively long skull with a narrow upper face [85]. In this regard, the shape effects associated with the peak marker in this interval, just downstream of the primary Prader-Willi genes, are suggestive, as they contrast a shorter, wider face with a longer, narrower one (see **Figs 5 and 6**, below).

**Fig 5.**
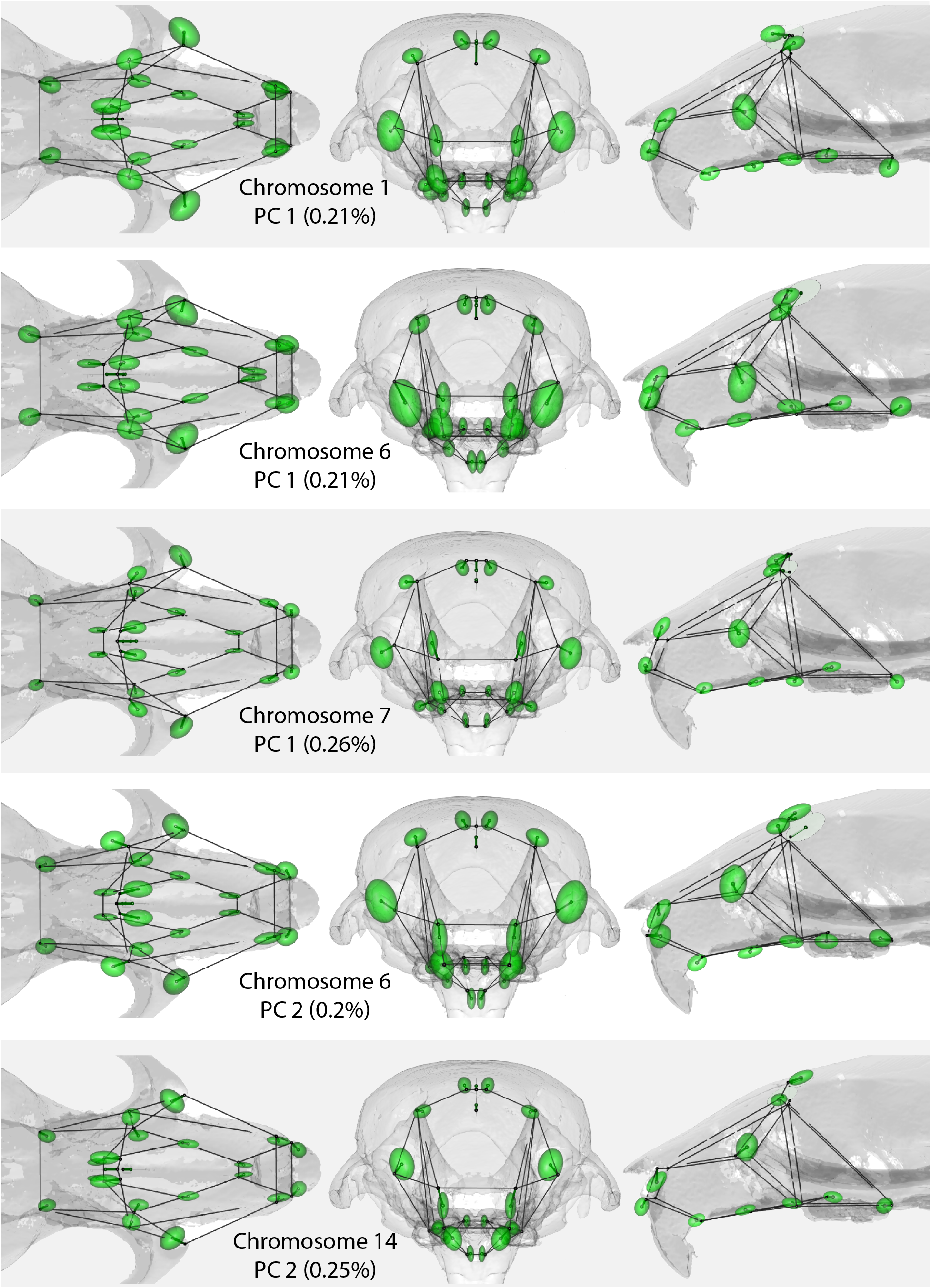
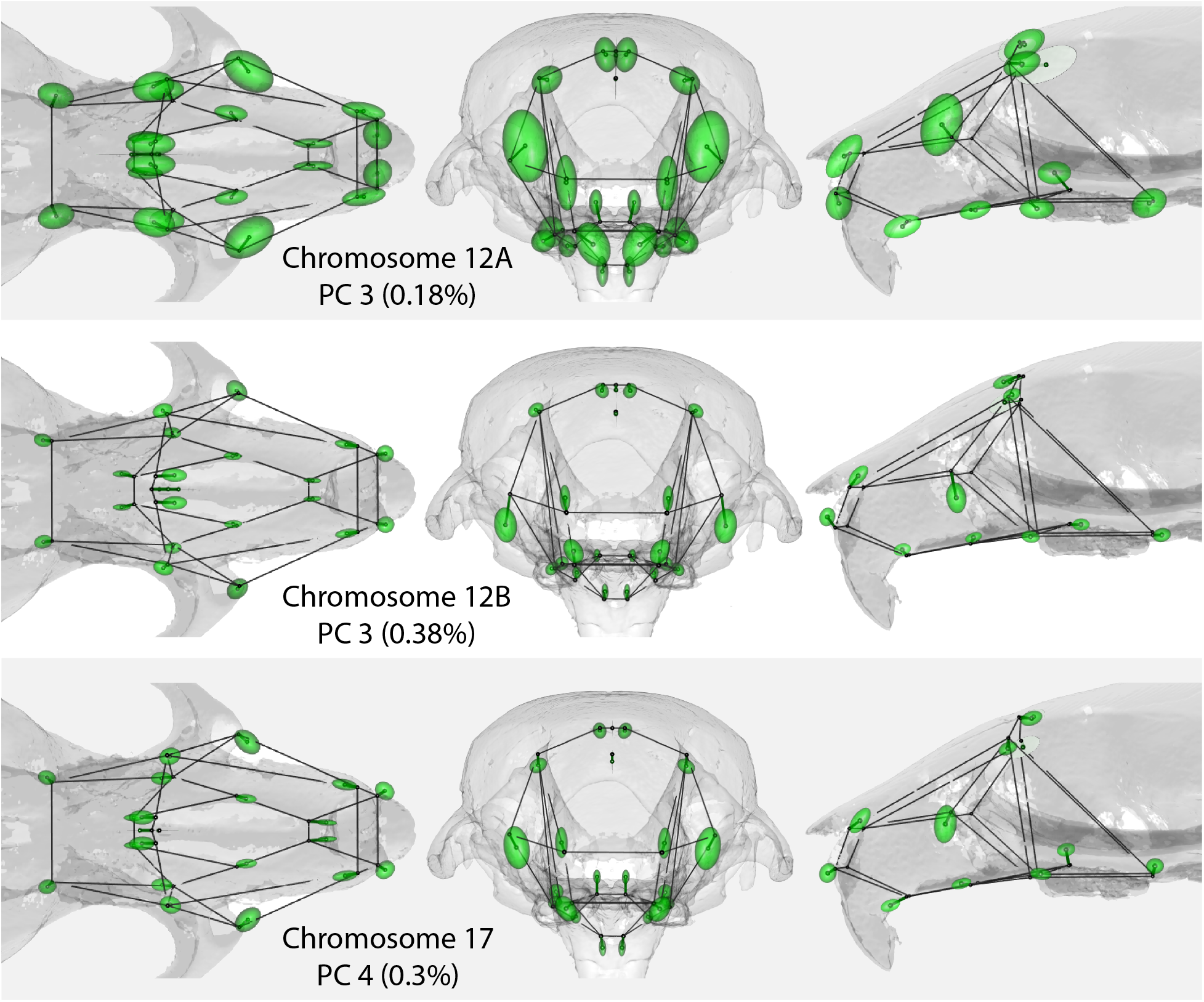
Superior (left), anterior (middle), and lateral (right) views of the (magnified) effect of allele substitution at the genome scan peak marker. Displacement vectors from wireframe to center of ellipsoids depict the mean effect. Ellipsoids enclose the approximate 80% posterior confidence boundaries.

**Fig 6.**
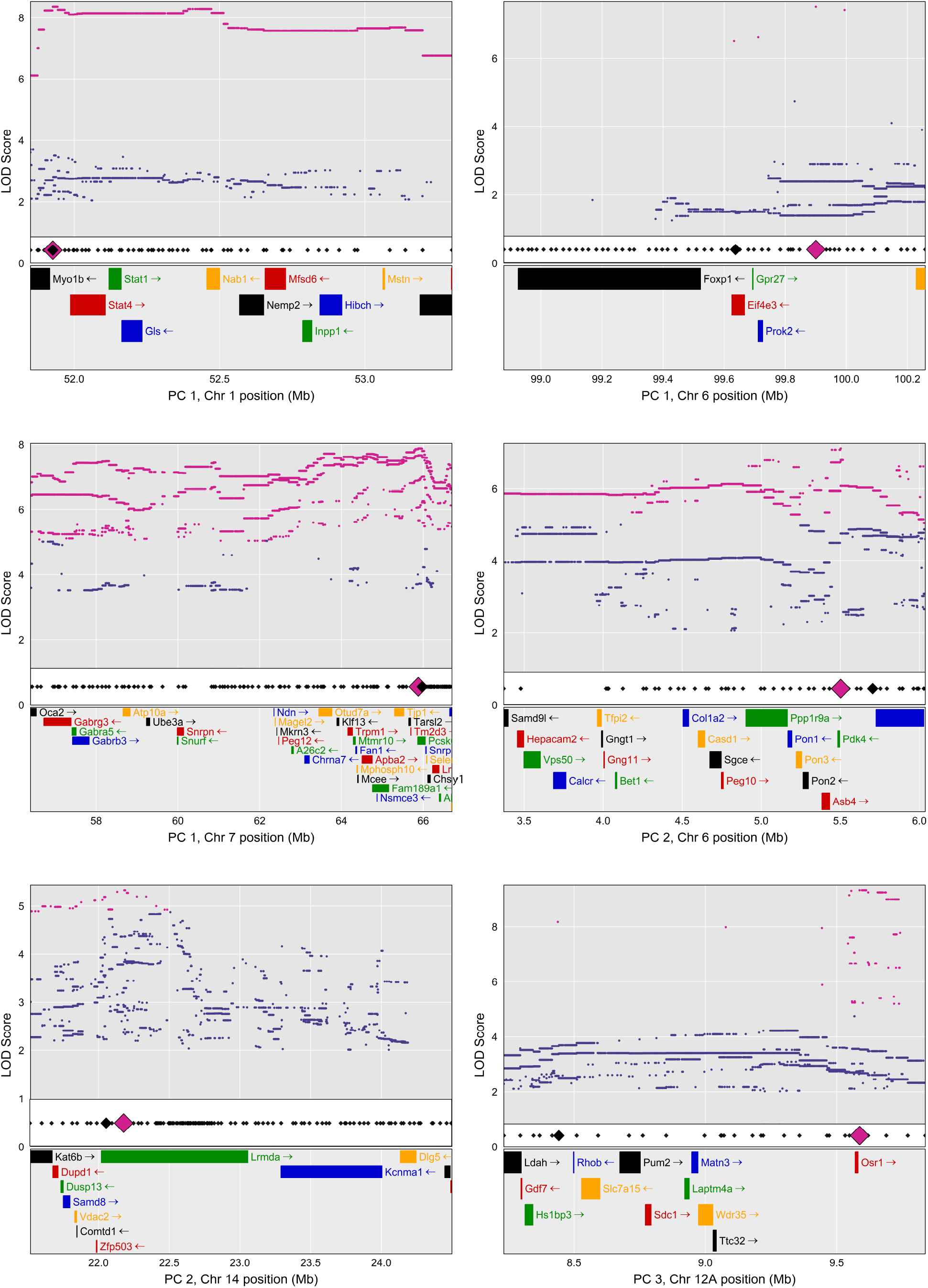
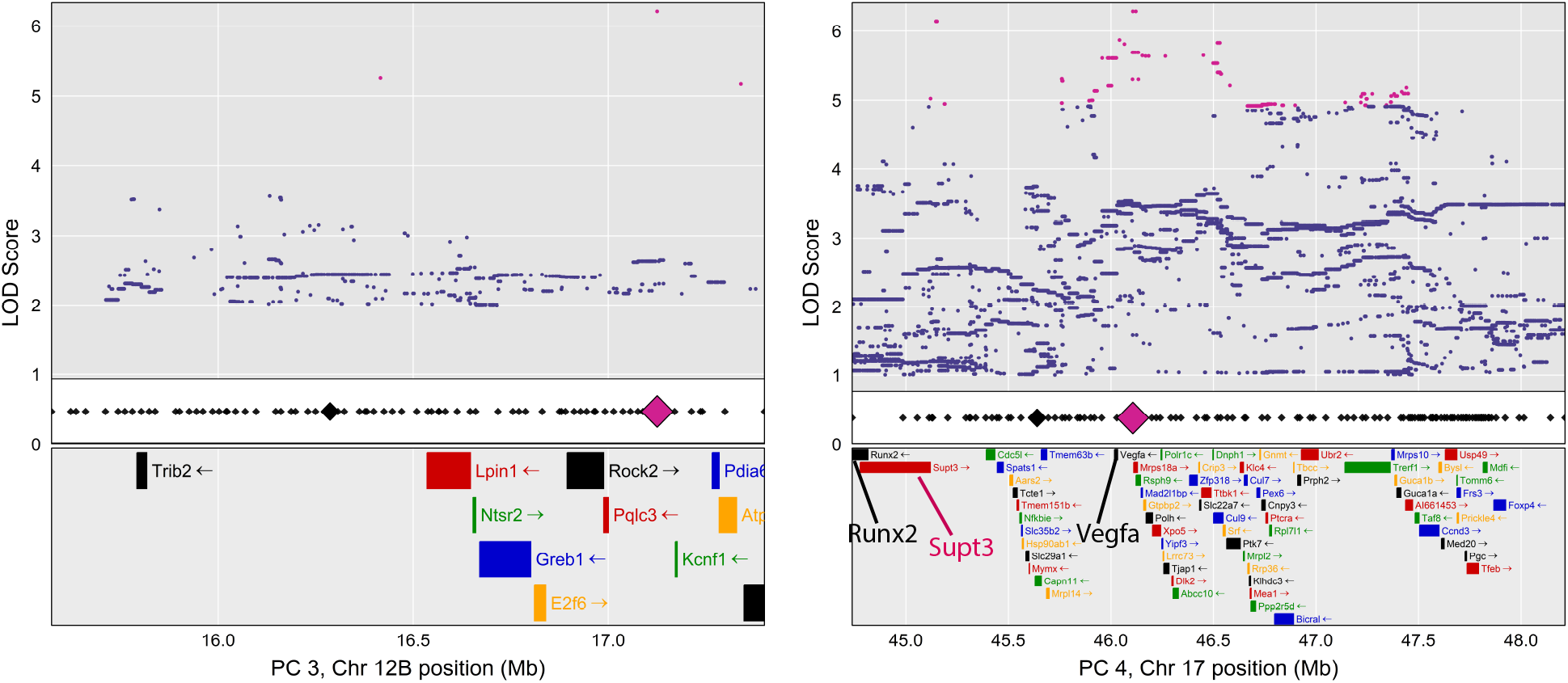
Association mapping results for PC 1-4 QTL intervals. Top panels: LOD scores for all CC SNP variants in the interval. Magenta points are locations where inferred bi-allelic SNP probabilities (based on strain distribution patterns in the CC) exceed the 95% LOD threshold. To control file size, low LOD scores points have been omitted from most plots. Middle panel: location of genotyped markers in the DO (small black diamonds), highlighting LOD peak locations for the founder genome scan (large black diamond) and SNP association scan (large magenta diamond). Bottom panel: named, protein coding genes for intervals. For legibility, we show the portion of the PC 2, Chromosome 14 map that contains both peak LOD scores and all SNPs exceeding the LOD threshold. For all plots, the complete gene list is provided in **S1 File**.

### Marker Effects on Shape

At the LOD peak marker of each PC 1-4 QTL, we collapsed the 8-state founder probabilities to 2-state SNP probabilities in order to estimate the shape effect of allelic substitution at that marker. To estimate effects on total shape (instead of univariate shape PCs), we employed a Bayesian mixed effects model for highly multivariate dependent observations [50, 86]. The results are plotted in **Fig 5**. In each figure, QTL effects are magnified by the scaling factor required to cause the QTL to account for 10% of phenotypic variance. These effects are shown as displacement vectors emanating from the mean configuration wireframe. The magnification ensures that even when the effect at a landmark is small, its direction and relative size (in comparison to effects at other landmarks) is apparent. We further exploited the Bayesian posterior to plot 80% confidence ellipsoids for the magnified displacements. The actual proportion of variance explained by the SNP, without magnification, is noted in the figure (and included in **Table 3**).

**Table 2.** Gene ontology enrichment for selected biological processes.

| Biological process | FDR $q$-value | GO annotated genes |
| --- | --- | --- |
| Animal organ morphogenesis | 1.42 e-3 | **PC1:** *Aldh1a3, Chsy1, Nab1, Stat1, Trpm1*. **PC2:** *Col1a2, Dlg5, Gngt1, Hesx1, Tnnc1, Wnt5a, Zmiz1*. **PC3:** *Gdf7, Matn3, Osr1, Sdc1*. **PC4:** *Ccdc66, Mdfi, Ptk7, Runx2, Srf, Vegfa*. |
| Growth | 2.25 e-3 | **PC1:** *Apba2, Mstn, Ndn, Nsmce3, Ube3a*. **PC2:** *Bap1, Colq, Il17rb, Mustn1, Sema3g, Tkt, Wnt5a, Zmiz1*. **PC3:** *Gdf7, Matn3, Osr1*. **PC4:** *Dnph1, Mymx, Ptk7, Srf, Taf8, Vegfa, Hsp90ab1*. |
| Skeletal system development | 1.29 e-2 | **PC1:** *Chsy1, Nab1, Foxp1, Lrrk1*. **PC2:** *Col1a2, Mustn1, Wnt5a*. **PC3:** *Matn3, Osr1, Runx2*. **PC4:** *Mdfi, Srf*. |
| Sensory organ development | 1.62 e-2 | **PC1:** *Aldh1a3, Gabra5, Gabrb3, Trpm1*. **PC2:** *Ccdc66, Gngt1, Hesx1, Wnt5a*. **PC3:** *Osr1*. **PC4:** *Prph2, Ptk7, Srf, Vegfa*. |
| Cartilage development | 2.4 e-2 | **PC1:** *Chsy1*. **PC2:** *Mustn1, Wnt5a*. **PC3:** *Matn3, Osr1*. **PC4:** *Runx2, Srf*. |
| Bone development | 2.54 e-2 | **PC1:** *Chsy1, Nab1, Lrrk1, Foxp1*. **PC3:** *Matn3*. **PC4:** *Runx2, Srf*. |

**Table 3.** Candidate genes.

| PC | Chr | Founder |  | Founder type | SNP |  | SNP_type | SNP % Var | Candidates |
| --- | --- | --- | --- | --- | --- | --- | --- | --- | --- |
|  |  | Peak (Mb) | Founder ID |  | Peak (Mb) | SNP ID |  |  |  |
| PC1 | 1 | 51.926616 | rs48415864 | intergenic | 51.926623 | rs254750887 | intergenic | 0.21 | *STAT1* |
| PC1 | 6 | 99.637221 | rs226005706 | intron | 99.900271 | rs31234872 | intergenic | 0.21 | *FOXP1* |
| PC1 | 7 | 65.970847 | rs33905059 | intron | 65.876674 | rs32405988 | intron | 0.26 | *CHSY1* |
| PC2 | 6 | 5.704731 | rs49620919 | intergenic | 5.502445 | rs32485785 | intergenic | 0.2 | NA |
| PC2 | 14 | 22.058402 | rs30739483 | intron | 22.182166 | rs30740474 | intron | 0.25 | *KAT6B* |
| PC3 | 12 | 8.44203 | rs48833407 | intergenic | 9.587123 | rs29212735 | intergenic | 0.18 | *OSR1*, *GDF7* |
| PC3 | 12 | 16.286802 | rs49949506 | intergenic | 17.125642 | SV 17121898: 17129386 | structural (deletion) | 0.38 | NA |
| PC4 | 17 | 45.639772 | rs107675960 | intron | 46.106749 | rs108721614 | upstream gene variant | 0.3 | *VEGFA*<br>*[RUNX2, SUPT3]* |

Candidate genes located within 0.25 Mb of founder or SNP QTL peaks. SNP % Var gives the percentage of variation explained by substitution of an additional reference allele at the founder (genome scan) peak.

### Biological Process Enrichment

We used the human annotations in the online Molecular Signatures Database [87] to query biological processes relevant to craniofacial variation are overrepresented in the PC 1-4 QTL intervals. Collectively, the intervals contain 243 named, protein coding genes (**S1 File**), 232 of which are annotated in the Entrez Gene library utilized by the Molecular Signatures Database. **Table 2** provides GO biological process enrichment highlights for these intervals, including FDR *q*-values adjusted for multiple testing (see **S2 File** for the top 100 *q*-value processes). Growth, morphogenesis, and skeletal system development are prominent biological process annotations for the facial shape QTL. The enrichment results suggest that no enriched biological process is uniquely associated with shape variation on a single PC. Rather, with the exception of bone development, each process in **Table 2** includes annotations to each PC.

To consider annotations specific to allometric variation, we queried biological process enrichment for PC 1 QTL alone (**S3 File**). Collectively, the three allometry QTL intervals were significantly enriched for genes implicated in bone development (FDR *q* = 0.025). Each of the intervals included at least one such gene: *Nab1* (chromosome 1) [88], *Foxp1* (chromosome 6) [89, 90], and *Lrrk1* and *Chsy1* (chromosome 7) [91–93]. The chromosome 1 interval also includes Myostatin (*Mstn, Gdf8*), a negative regulator of muscle growth. Inactivation of *Mstn* has been shown to greatly increase facial muscle mass, with corresponding size decreases and shape changes throughout the bony face and vault [94, 95].

### Association Mapping

Finally, to develop a finer-grained impression of potential causal loci, we performed SNP association mapping within each QTL interval. We imputed a dense map of SNP probabilities to the sample by inferring their bi-allelic (major/minor) probabilities based on (a) founder probabilities at flanking, genotyped markers and (b) major/minor strain distribution patterns in the CC founder variants **(Fig 6)**.

We then queried whether any genes within 0.25 Mb of the SNP or genome scan LOD peaks are known to be associated with craniofacial development or phenotypes **(Table 3)**. As with human facial shape GWAS [42], LOD peaks tended to be located in non-coding regions. While the focal genes we identified all have known roles in skeletal or craniofacial development, they also tend to be developmentally ubiquitous. *Stat1* (PC1, chr. 1) is implicated in bone development and remodeling throughout life [96–98]. Overexpression of *Foxp1* (PC1, chr. 6) can attenuate osteoblast differentiation, chondrocyte hypertrophy, osteoclastogenesis, and bone resorption, while *Foxp1* deficiency can accelerate ossification [89, 90]. In humans, some *Foxp1* mutations produce a facial phenotype that includes broad forehead and short nose [99]. Chondrogenesis is generally disrupted in *Chsy1* null mouse mutants (PC1, chr. 7), and in humans, *Chsy1* mutations can manifest in micrognathia and short stature [92, 100]. *Kat6b* (PC2, Chr. 14) activates the transcription factor *Runx2*, which is often referred to as a master regulator of bone formation [101–103]. For the PC3 QTL at 8.23 – 9.84 Mb on chromosome 12, the founder and SNP LOD peaks reside at opposite ends of the QTL interval, but each is very close to an important craniofacial development gene: *Gdf7* (*Bmp12*) contributes to patterning of the jaw joint and is expressed in the developing incisor pulp and periodontium [104–106]; *Osr1* is broadly expressed in the embryonic and developing skull, with overexpression resulting in intramembranous ossification deficiencies, increased chondrocyte differentiation, and incomplete suture closure, while under-expression may contribute to cleft palate phenotype [107–109]. *Osr1* has also previously been identified as a candidate in human facial shape GWAS [23]. Finally, the SNP LOD peak for PC4, chromosome 17 implicates *Vegfa*, which is critical to endochondral and intramembranous ossification through its role in angiogenesis [110]. A distinct SNP LOD peak of nearly equal magnitude is located approximately 1 Mb upstream (at rs47194347), very close to *Supt3* (human ortholog *Supt3h*) and *Runx2*. In addition to the central role *Runx2* has in bone formation and development [102], both genes are human facial shape GWAS candidates [15, 22].

While GP maps for traits influenced by many genes may be quite intricate and layered, additive effect QTL analyses remain an indispensable means to identify loci that contribute to phenotypic variation. Our investigation advances understanding of the genetics of facial skeletal diversity in several ways. The 37 QTL identified herein substantially expand the small but growing facial shape QTL library. Compared to similar prior studies, the number of QTL is relatively large and their confidence intervals relatively narrow. These results affirm the value of larger samples, sophisticated animal models like the outcrossed population utilized here, and high-resolution genotyping for identification and localization of QTL. Synthesizing our results with prior insights from developmental biology and medical genetics provides a richer picture of the sources of facial skeletal diversity. We found a relative abundance of skeletal and craniofacial development genes in QTL intervals for the largest shape PCs. Moreover, these genes tend to be located very close to QTL peaks. Thus, our results suggest that genes essential to building a face are often also the genes that cause faces to vary. Finally, QTL effect sizes were small. In summary, our results are consistent with a model of facial diversity that is influenced by key genes in skeletal and facial development and, simultaneously, highly polygenic.

## Materials and methods

### Diversity Outbred mouse population

The DO (Jackson Laboratory, www.jax.org) are derived from the eight inbred founder lines that contribute to the CC [43, 44], which include three mouse subspecies, and both wild-derived and laboratory strains [46]. Continued random outcrossing of the DO has produced a population with relatively high genetic diversity (∼45 million SNPs segregating in the CC founding populations [43]) and increasingly fine mapping resolution in advancing generations.

In combination, these features increase the potential to identify multiple QTL for complex traits [31]. On average, the genomes of the eight CC founders should be equally represented in the DNA of any DO animal. Sequenced genomes for the CC make it possible to characterize and interpret genetic variants in the DO according to 8-state founder identity and as bi-allelic (major/minor allele) SNP variation. In addition, statistical packages designed for analysis of the DO offer features to impute variants at each typed SNP in the CC founders, using the coarser DO marker map as a scaffold to infer a high-density SNP map.

### Computing

All analysis was conducted in R [111]. Customized data processing and statistical analysis functions for mapping experiments in DO samples were originally published in the DOQTL package [49]. These routines are now incorporated into the qlt2 package [48], with additional processing and genotype diagnostic tools provided at https://kbroman.org/qtl2/.

### Images, landmark data

Our DO sample consists of *n=*1147 male and female adult mice (after exclusion of some records; see *Genotypes*, below) from seven DO generations (**Table 4**). We obtained µCT images of the mouse skulls in the 3D Morphometrics Centre at the University of Calgary on a Scanco vivaCT40 scanner (Scanco Medical, Brüttisellen, Switzerland) at 0.035 mm voxel dimensions at 55kV and 72-145 µA. On each skull, one of us (WL) collected 54 3D landmark coordinates from minimum threshold-defined bone surface with ANALYZE 3D (www.mayo.edu/bir/). Here, we analyzed the 22-landmark subset of the full configuration—a 66 coordinate dimension vector—that captures the shape of the face (**Fig 1**).

**Table 4.** DO sample composition.

|  | **DO generation** |  |  |  |  |  |  |
| --- | --- | --- | --- | --- | --- | --- | --- |
|  | <u>**9**</u> | <u>**10**</u> | <u>**15**</u> | <u>**19**</u> | <u>**21**</u> | <u>**23**</u> | <u>**27**</u> |
| **Female** | 91 | 221 | 80 | 55 | 80 | 97 | 75 |
| **Male** | 0 | 144 | 0 | 58 | 77 | 99 | 70 |
| **MUGA Array** | Mega | Mega | Mega | Giga | Giga | Giga | Giga |

Overview of the DO sample by generation, sex, and genotyping array.

### Ethics Statement

No animals were sacrificed for this study. Specimens were raised in several labs for unrelated experiments under approval and conduced in accordance with the guidelines set forth by the Institutional Animal Care and Use Committee in accordance with protocols #11-299 (University of North Carolina Chapel Hill), #99066 and #15026 (Jackson Labs), and #08-0150-3 (Scripps Research Institute). Receipt of animal carcasses and micro-CT (μCT) imaging of mouse skulls were undertaken in accordance with approved IACUC protocol #AC18-0026 at the University of Calgary.

### Shape data

Landmark configurations were symmetrically superimposed by Generalized Procrustes Analysis (GPA) [112] using the Morpho package [113]. GPA is a multi-step procedure that removes location and centroid size differences between configurations, then (iteratively) rotates each configuration to minimize its squared distance from the sample mean. We chose a superimposition that decomposes shape variation into its symmetric and asymmetric components [114, 115], and stored the symmetric configurations for analysis. Working with symmetric, superimposed data reduced morphological degrees of freedom from 66 landmark coordinate dimensions to 30 shape dimensions [115].

### Genotypes

For all animals, DNA was extracted from ear or tail clippings. Genotyping was performed by the Geneseek operations of Neogen Corp. (Lincoln, NE). Mice from generations 9, 10, and 15 were genotyped using the MegaMUGA genotyping array (77,808 markers); mice from generations 19, 21, 23, and 27 were genotyped using the larger GigaMUGA array (143,259 markers) [116]. The two arrays share 58,907 markers in common. We restricted the genotype data to these shared SNPs [116].

In addition, we excluded 2,022 markers from the center of chromosome 2 (40 - 140 Mb). WSB alleles are highly overrepresented in this genomic region in earlier DO generations. This overrepresentation has been attributed to female meiotic drive favoring WSB at the *R2d2* locus [117]. Between DO generations 12 and 21, Jackson Labs systematically purged the meiotic drive locus and linked effects from the DO [45]. Our samples closely track this history, with WSB allele frequencies in the middle of chromosome 2 exceeding 60% in generation 9 and falling to 0% by generation 23 (**S2 Fig**). If unaddressed, this directional shift in allele frequencies can confound statistical analysis of genotype-phenotype associations [118] with potentially false positive associations between WSB chromosome 2 haplotypes and directional changes in phenotype over DO generations. After removing this genomic region from consideration, the final SNP dataset consists of 56,885 markers common to the MegaMUGA and GigaMUGA arrays, with a mean spacing of 48 kb.

The qtl2 package estimates genotype probabilities, including imputation of sporadically missing genotypes, with a multipoint Hidden Markov Model (HMM) on the observed marker genotypes. From these initial genotype probabilities, we derived three probabilistic representations of genetic state for each observation at each marker: (1) for genome scans, founder probabilities, a vector encoding the probability of genetic contribution from each of the eight CC founders to the diplotype state; (2) for association mapping, SNP probabilities, collapsing the eight founder probabilities to a two-state, major/minor allele strain distribution; and (3) for mapping multivariate shape effects, additive SNP dosage, twice the estimated major SNP probability (maximum additive SNP dosage = 2).

We identified problematic genetic samples using the DO genotype diagnostics tools available at https://kbroman.org/qtl2/ [119]. A total of 11 observations were excluded due to excessive missing marker data, which we defined as no marker information at >7.5% of markers. Three probable duplicate sets of genetic data (>99% of markers identical) were identified. Because the correct genotype-phenotype matches could not be established by other evidence, all six animals were removed from the sample. Finally, we excluded seven female samples with low X chromosome array intensity values. These samples are potentially XO [45], which can affect facial form and body size [120, 121]. The exclusions described above resulted in a final, retained sample of *n* = 1147 animals.

### Genome scan and association mapping

We mapped QTL for facial shape and facial shape allometry. To identify QTL, we implemented the linear mixed effects regression model in qtl2. The qtl2 genome scan is designed to estimate marker effects on one or more univariate observations, but not covariance among those dimensions. We therefore decomposed the superimposed coordinates into PCs, treating the specimen scores on each PC as measured traits. For allometry, we treated common allometric component scores [61] as measured traits. Prior to deriving PCs and allometry vectors, we centered each observation on its generation mean in order to focus on shared, within-generation covariance patterns [122, p. 295].

At each marker, we fit a mixed model with fixed effect coefficients for sex, age-at-sacrifice, and the additive effect of each CC founder haplotype, a random effect of kinship, and an error term [49]. Evidence for a QTL was evaluated based on marker LOD scores, likelihood ratios comparing the residuals of the marker model to a null model excluding the marker’s effect [51]. We consider the evidence for a marker-phenotype association to be statistically significant where the LOD score exceeds the 95% quantile of genome-wide maximum LOD scores computed from 1000 random permutations of genotype-phenotype associations [123]. We use a 1.5 LOD-drop rule to define the boundaries of QTL support intervals [124], and allow for multiple QTL peaks on a single chromosome.

The random effect of kinship depends upon pairwise proportions of shared alleles among observations, encoded in a kinship matrix. Loss of power to detect QTL can arise when the marker of interest and linked genes are included in the kinship matrix of the null model. To avoid this problem, we compute kinship matrices using a leave-one-chromosome-out (LOCO) approach [52–54].

Genome scans, QTL support intervals, and proportions of shape variation attributable to founder-level kinship (heritability) were estimated for all PCs explaining more than 1% of symmetric shape variation. However, we limit more in-depth analysis to the five PCs with reasonably large signal-to-noise ratios (**S2 Fig**). This was assessed using the modified “noise floor” detection strategy developed by Marroig and colleagues in [56].

In addition, within the PC 1-5 QTL intervals, we used a form of merge analysis to perform genome-wide association mapping [49, 125, 126]. For each CC founder variant located between a flanking pair of genotyped markers, the vector of founder allele probabilities is assumed to be the average of the founder probabilities at those flanking markers. The bi-allelic SNP probabilities were computed from these founder probabilities and the major/minor allele strain distribution pattern in the CC. Analysis was again via mixed effects regression model, with the imputed SNP data as genotype predictors.

### Enrichment analysis and candidate genes

Modern GWAS and QTL studies often attempt to contextualize QTL and candidate gene results post hoc via GO enrichment analysis. Given a list of genes near statistically significant markers, a GO analysis calculates the probability that a random set of genes of the same size would contain as many genes under a common annotation term, typically after a family-wise error rate or FDR adjustment for multiple testing. The outcome of enrichment analysis is a list of GO terms (or in the absence of any enrichment, no list), which provides a means to consider whether certain biological processes, cellular components, or molecular functions are over-represented in the list of genes [127].

For protein coding genes in the PC 1-5 QTL support intervals and for allometry QTL, we used the Molecular Signatures Database [87] online tool and Entrez Gene library [128] to identify biological process GO term enrichment [129]. The database application implements a hypergeometric test of statistical significance, quantifying the probability of sampling, without replacement, as many (*k*) or more genes annotated to a GO term, given the total number of genes annotated to that term (*K*) and the number of genes in the QTL and Entrez gene sets (*n* and *N*, respectively). For each ontology term, after adjustment for multiple testing, we required a FDR *q*-value < 0.05 to conclude that there was substantial evidence for GO biological process enrichment. We also inspected the association mapping results in the +/- 0.25 Mb region centered at each LOD peak for genes with convincing evidence of a role in skeletal, craniofacial, or dental development or disease. For most QTL, LOD peaks for the genome scan and association mapping results are virtually coincident. Where they differ substantially, we examine the region around both peaks.

### Marker effects on shape

We considered a significant association between a marker and facial shape along a PC axis to be evidence of some effect on the multivariate (undecomposed) shape coordinate data. In order to obtain an estimate of the full shape effect, we fit an additional mixed model for the maximum LOD score marker. Fitting a mixed model for highly multivariate data presents a significant computational challenge because the number of parameters that must be estimated for the random effects covariance matrix scales quadratically with the number of traits (here, 465 parameters for 30 free variables). To address this challenge, we estimated mixed model covariance matrices using a Bayesian sparse factorization approach [50, 86, 130]. After a 1,000,000 iteration burn-in, we generated 1,000,000 realizations from a single Markov chain, thinning at a rate of 1,000 to store 1,000 posterior samples for inference. Though the time needed to fit such a model makes it impractical for a genome scan on a dense array, it is a useful strategy for disentangling kinship from the effect on multivariate outcomes at a small number of peak markers. For these regressions, the marker effect predictor was twice the major SNP probability. Thus, the multivariate mixed model we implemented estimates an average effect of allele substitution, a standard quantity in quantitative genetics [131]. In addition, for each regression, we estimated the proportion of shape variation explained using a version of the coefficient of determination (*R*^2^) that has been adapted for mixed model parameters [132].

When plotting shape effects, we magnify the allele substitution coefficients by the scaling factor needed to cause the QTL to explain 5% of phenotypic variance. This magnification is applied both to the mean effect, in order to depict the mean displacement vector, and to the posterior distribution of effects, in order to depict confidence ellipsoids. Ellipsoids were estimated with the 3D ellipse function in the R package rgl [133]. While it is more typical to use a 95% support boundary to evaluate effect uncertainty, with shape data, morphological contrasts are best understood over the configuration as a whole [134]. The 80% ellipsoids are meant to encourage this perspective.

## Acknowledgements

Thanks to Xiang Zhao for help with preparation of samples and David Eby for helpful intervention with the online Molecular Signatures Database. We thank Kunjie Hua, Liyang Zhao and Kuo-Chen Jung for assistance with UNC mouse experiments. Charles Farber is gratefully acknowledged for assistance with processing of UNC mouse carcasses. All UNC mouse procedures were approved and guided by the appropriate institutional animal ethics committees, including the University of North Carolina at Chapel Hill.

## Supporting Information

**S1 Fig. Eigenvalue decay**. Percentage of total variation explained for each shape principal component. We performed genome scans for all PCs explaining more than 1% of shape variation (red point identifies cutoff at PC 20) and subjected the first five PCs (green point identifies PC 5 floor) to in-depth analysis.

**S2 Fig. Change in chromosome 2 WSB allele frequencies over generations.** Sample-wide WSB allele frequency on chromosome 2 by DO generation. The high WSB allele frequency in earlier generations reflects meiotic drive for WSB at the *R2d2* locus. Declining frequencies in later generations reflect the systematic purge of WSB alleles from the DO population.

**S1 Table.** LOD peaks and support intervals for all QTL on PCs 1-20.

**S2 Table.** Full array of LOCO mean heritabilities over PCs 1-20.

**S1 File. Genes, PCs 1-5.** All named, protein-coding genes in the PC 1-5 QTL intervals.

**S2 File. Top GO annotations, PCs 1-5.** Top 100 FDR q-value GO terms for the PC 1-5 QTL interval gene set (Molecular Signatures Database output file).

**S3 File. Top GO annotations, PC 1.** For the PC1 QTL interval gene set, GO terms with FDR q-value < 0.05 (Molecular Signatures Database output file).

